# Identification and characterisation of variable rhythms using CosinorPy

**DOI:** 10.1101/2022.07.04.498691

**Authors:** Miha Moškon

## Abstract

**Background and objective:** Analysis of rhythmic data has become an important aspect in biological and medical science, as well as other fields of science. Parametric methods based on trigonometric regression reflect several advantages in comparison to their alternatives. Software packages for parametric analysis of rhythmic data are mostly based on a single-component cosinor model, which is not able to describe asymmetric rhythms that might reflect multiple peaks per period. Moreover, a basic cosinor model is unable to describe rhythmicity that is changed with time. Here, we present some important extensions of the recently developed CosinorPy Python package to address these gaps.

**Methods:** The extended package CosinorPy provides the functionalities to perform a detailed individual or comparative analysis of rhythms reflecting (1) multiple asymmetric peaks per period, (2) forced, damped, or sustained rhythms, and (3) shift of the midline statistics of rhythm over time. In all these cases the package is able to assess the (differential) rhythmicity parameters, evaluate their significance and confidence intervals, and provide a set of publication-ready figures.

**Results:** We demonstrate the package in some typical scenarios that incorporate different types of rhythmic dynamics, such as asymmetric, damped, and forced rhythms. We show that the proposed implementation of a generalised cosinor model is capable of reducing the error of estimated rhythmicity parameters in cases that tend to be problematic for alternative models. The implementation of the presented package is available together with scripts to reproduce the reported results at https://github.com/mmoskon/CosinorPy.

**Conclusion:** According to our knowledge, CosinorPy currently presents the only implementation that is able to cover all the above scenarios using a parametric model with vast applications in biology and medicine as well as in other scientific domains.

## 1. Introduction

Identification and analysis of rhythmic data, for example circadian data, have become important in different aspects of science, especially in the context of biology and medicine. Several non-parametric approaches aimed at such analyses have been reported in recent years (for examples, see [1, 2, 3, 4, 5]). However, parametric approaches mostly based on trigonometric regression still reflect several advantages over nonparametric approaches in many cases. Parametric methods can be applied even in cases when we are dealing with unevenly spaced and/or unbalanced data [6, 7, 8, 9, 10]. Rhythmicity parameters, such as amplitudes and acrophases, can be derived directly from the coefficients of a regression model, which is not the case for many non-parametric methods [2]. Parametric methods can be used to assess the significance of each rhythmicity parameter that is different from zero [11] as well as to compare rhythmicity parameters among different measurements and to assess their differential rhythmicity using easy-to-interpret p-values [9, 12, 13, 14]. Parametric methods can be adapted to different situations and can be used to assess the population mean response [9, 11, 12, 15] as well in a combination with different count models for the analysis of count data (for example, see [16]). Nonparametric methods are mostly aimed towards the analysis of omics, e.g., transcriptomics data. On the other hand, parametric approaches have much wider applicability. They can be applied to different domains even outside biology and medicine (for example, see [17]).

Recent parametric approaches for identification and analysis of rhythmic data are mostly based on a cosinor model [11], and include ECHO (extended circadian harmonic oscillator) [18], COMPARERHYTHMS [19], CircaCompare [14] and CosinorPy [9]. ECHO is based on a generalised cosinor model which is able to describe sustained (as in the case of a basic cosinor model), damped, or forced rhythms. However, a threshold used for the classification of a measurement in any of these three groups is arbitrarily defined and might depend on the noise in the data [18]. Moreover, ECHO is limited to a single-component cosinor model, which means it can only describe the rhythms reflecting a single and a symmetric peak per period. COMPARERHYTHMS provides a differential rhythmicity analysis pipeline based on the basic single component cosinor model. CircaCompare presents a method also focused on the evaluation of differential rhythmicity and is based on a non-linear regression in combination with the cosinor model [14]. It can account for attenuation or dampening of rhythm amplitudes over time, however it is again limited to a single-component model.

Recently, CosinorPy, a Python package for the identification and characterisation of rhythmic behaviour has been reported [9]. This package has already been used in different studies ranging from a the analysis of ‘circadian reprogramming’ in cell lines using qPCR data [20] to predicting sleep quality metrics [21] and chronic pain predictions [22]. CosinorPy implements single- as well as multicomponent cosinor models, which can be used in a combination with the population as well as count models. Moreover, the package can be used to assess the differential rhythmicity among pairs of measurements. However, if a user requires evaluation of more accurate parameter statistics, such as the significance of rhythm amplitudes and the significance of change between acrophases in pairs of measurements, a single-component cosinor had to be used in the basic implementation. In this model, the rhythmicity parameters and their corresponding p-values can be analytically evaluated using the delta method, as introduced in [23] and adapted in [9]. However, when using a multicomponent cosinor model, the rhythmicity parameters cannot be calculated directly from the regression coefficients, so alternative methods must be used. Their evaluation can be performed numerically from the fitted curves [9]. Here, we describe extensions to the multicomponent cosinor model that allow us to evaluate (1) the p-values of rhythmicity parameters in a multicomponent cosinor model, and (2) the p-values describing the significance of changes in the rhythmicity parameters in pairs of measurements using multicomponent cosinor models. The methodology behind these extensions is based on bootstrapping of confidence intervals and p-values of selected rhythmicity parameters. This increases the computation time required to obtain the results, but provides much richer statistics.

Furthermore, we extend the cosinor model to a generalised cosinor model that can account for the accumulation or reduction of the midline statistics of rhythm (MESOR) and to describe the damping or amplification of rhythm amplitudes over time. This is especially vital for longer time series (relative to the rhythmic period) during which the amplitude and MESOR are likely to change due to different factors [18, 24]. We extend both a single-component and a multicomponent cosinor model to its generalised version. The rhythmicity parameters, their corresponding confidence intervals, and p-values can be estimated directly from the regression coefficients in a single-component model. In a multi-component model, bootstrapping is used in a similar way as in a basic multi-component cosinor model. Contrary to ECHO, our implementation can yield a p-value describing the significance of damping or amplification of an amplitude over time and can be used even when data exhibit multiple asymmetric peaks within a rhythmic period. Furthermore, the proposed implementation can be used to evaluate the comparative rhythmicity of pairs of measurements to evaluate changes in rhythmicity parameters, such as the significance of the acrophase shift, the amplitude change, the change in the amplification coefficient or the change in the linear component.

The extended version of the CosinorPy package thus provides the functionalities to perform an individual or comparative analysis of rhythmic behaviour reflecting (1) multiple asymmetric peaks per period, (2) forced, damped, or sustained rhythms, and (3) accumulation or reduction of MESOR with time. To the best of our knowledge, it presents the only implementation that can cover all these scenarios using a parametric model with vast applications in biology and medicine, as well as in other scientific domains. The functionalities reported here are available in the extended CosinorPy package (version 2.1), which can be installed by `pip install CosinorPy==2.1`. All main functionalities of the package can be accessed through easy-to-use wrapper functions, as demonstrated in a series of interactive Python notebooks (IPYNB) available at https://github.com/mmoskon/CosinorPy.

## 2. Methods

### 2.1. From a single- to a multi-component cosinor model

A multicomponent cosinor model can be described by the following equation:

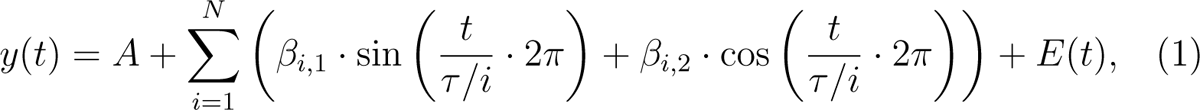

where *t* corresponds to the observed time points in the time series, *E*(*t*) is the error term, *τ* is the assumed rhythmicity period, *N* is the number of components in the model, and *A*, *β*_1,1_, *β*_1,2_, …, *β*_*N*,1_, *β*_*N*,2_ are the regression coefficients. In these, *A* describes the MESOR (midline statistics of rhythm) and *β_i,j_* the amplitude of a trigonometric component *i, j* [11].

If the period is known, this model can be transformed into a linear form, which simplifies the regression process [9, 11]. The linear regression behind the cosinor model is used to determine the values of the regression coefficients. If we assume that these parameters are distributed according to a t distribution, their confidence intervals as well as their p-values can be estimated using the t test. Using the values of regression coefficients, the rhythmicity parameters can be calculated analytically in a one-component cosinor model, which is not possible when the number of components is increased. On the other hand, the rhythmicity parameters of a multicomponent cosinor model can be evaluated from the fitted curve, as described in [9]. This can be combined with bootstrap resampling to evaluate the confidence intervals and p-values of rhythmicity parameters in a multicomponent cosinor model.

CosinorPy2 implements bootstrapping as a means of assessing the confidence intervals as well as the significance of a rhythmicity parameter. The observed measurements are resampled with repetitions (bootstrap samples), and a cosinor model is constructed from each of the bootstrap samples. The rhythmicity parameters are evaluated against these models to construct their bootstrap distributions. These are then used to evaluate the significance and confidence intervals of each of the observed rhythmicity parameters.

In addition, bootstrapping can also be applied to the evaluation of the significance of differential rhythmicity between two datasets. In a multicomponent cosinor model, this can be performed by evaluating confidence intervals of mean differences (for example, amplitudes) and assessing their corresponding p-values, with the null hypothesis being that the means are equal.

Another benefit of using bootstrapping to assess the distribution of rhythmicity parameters is that an arbitrary metric that can be numerically evaluated on the fitted curve can be analysed. For example, one could also opt to analyse the asymmetry of peaks, centre of gravity of the fitted curve, etc. Even though this would require manipulation of the source code of the package, the extensions would be straightforward, since only the function used to evaluate the rhythmicity parameters on a fitted curve would require a minor modification.

### 2.2. From a cosinor to a generalised cosinor model

A generalised single-component cosinor model can be described as follows:

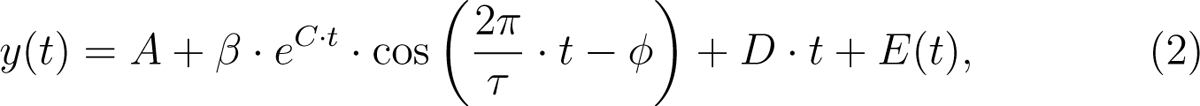

where *A* is the MESOR, *β* is the rhythmicity amplitude, *C* is the amplification coefficient, *D* is the linear component, *phi* is the acrophase, *τ* is the period, *t* is the time, and *E*(*t*) is the error term. The amplification coefficient defines whether a rhythm is sustained (*C* = 0), forced (*C* > 0), or damped with time (*C* < 0). This model cannot be expressed in a linear form. To estimate its coefficients, a nonlinear regression must be performed [9, 14]. The p-values of the individual coefficients and their corresponding confidence intervals can be evaluated using the covariance matrix obtained within the regression process.

A generalised cosinor model can be used to describe rhythms that are forced or damped over time in a manner similar to ECHO (extended circadian harmonic oscillator) [18]. In addition, the proposed generalised cosinor model can also describe linear trends within the data, namely, the accumulation or reduction of MESOR. Furthermore, the proposed model can be extended to perform a comparative analysis of the rhythmicity parameters between two datasets

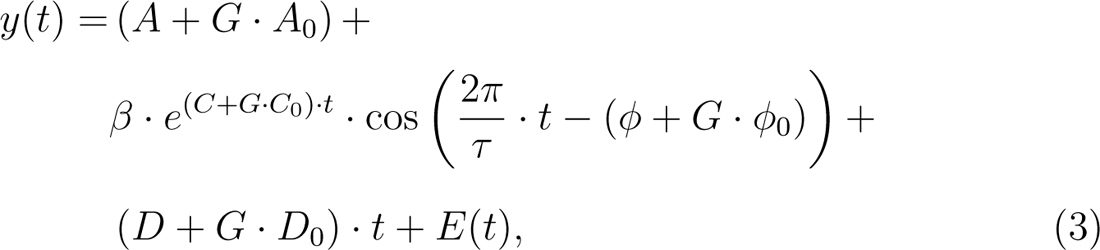

where *G* defines the dataset and is equal to 0 if a measurement belongs to the first dataset and 1 if it belongs to the second dataset. The coefficients *A*_0_, *B*_0_, *C*_0_, *D*_0_ and *ϕ*_0_ define the differences between the datasets. Their corresponding p-values can be used to assess whether the differences between the datasets are significant for each rhythmicity parameter.

A generalised cosinor model can be extended to a multicomponent generalised cosinor model:

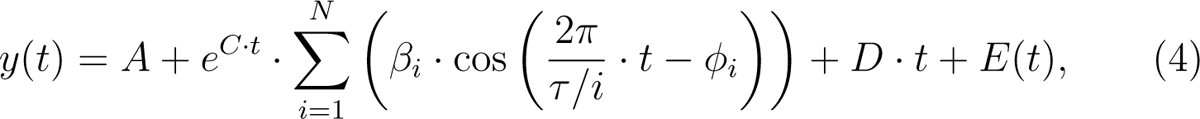

where *β_i_* represents the amplitude of component *i* and *ϕ_i_* its acrophase. As in the case of a basic multicomponent cosinor model, bootstrap resampling can be applied to evaluate the confidence intervals and p-values of the rhythmicity parameters in a generalised multicomponent cosinor model (see Section 3.1).

In addition, the differential rhythmicity statistic between two datasets can be evaluated similarly as in a basic model, namely by using bootstrap for the evaluation of amplitude changes and acrophase shifts. However, significance in the change of MESOR (*A*), amplification coefficient (*C*) and linear component (*D*) can be assessed directly from the regression coefficients and their covariance matrices (see Section 3.2).

## 3. Results

### 3.1. Analysing different types of rhythmic data

We used synthetically generated data to demonstrate the functionalities of the extended CosinorPy package on some typical scenarios and their combinations. Namely, we generated data reflecting (1) symmetric and asymmetric rhythms with single or multiple peaks per period, (2) damped, sustained, and forced rhythms, and (3) with or without the accumulation of a linear component around which rhythmicity occurs (MESOR). The generation process is summarised in Table 1 and available together with the presented data analysis pipeline as interactive Python Jupyter notebook at https://github.com/mmoskon/CosinorPy/blob/master/demo_independent_nonlin.ipynb. First, we analysed the generated data using a basic multi-component cosinor model as described in [9]. CosinorPy was used to identify the optimal number of components in a model to describe the given data as accurately as possible. The results of the fitting process are visualised in Figure 1 and included as supplementary Table 1. A more detailed analysis of the results obtained with multi-component cosinor models was performed using bootstrap as described in Section 2 (see Supplementary Table 2). However, as seen from Figure 1, this model was unable to accurately describe forced or damped rhythms (see `sym_damped`, `sym_forced`, `asym_damped`, `asym_forced`) and/or the accumulation of a linear component around which rhythmicity occurs (see `sym_lin_comp`, `asym_lin_comp`).

**Table 1:** Parameters for synthetically generated data. Rhythmicity periods of the generated data were set to 24 hours in all scenarios. Sampling period was set to 2 hours and noise level to 30%. Each scenario had three replicates. Abbreviations and symbols: *N* – number of components in a cosinor model, *ϕ* – acrophase; amplif. – amplification; lin. – linear component

| test | $N$ | sampling time | $\phi$ | amplif. | lin. |
| --- | --- | --- | --- | --- | --- |
| sym | 1 | 48 h | $\pi$ | 0 | 0 |
| sym_lin_comp | 1 | 48 h | 0 | 0 | 0.1 |
| asym | 3 | 48 h | $\pi$ | 0 | 0 |
| asym_lin_comp | 3 | 48 h | 0 | 0 | 0.1 |
| sym_damped | 1 | 72 h | 0 | -0.04 | 0 |
| sym_forced | 1 | 72 h | $\pi$ | 0.04 | 0 |
| asym_damped | 3 | 72 h | 0 | -0.04 | 0 |
| asym_forced | 3 | 72 h | $\pi$ | 0.04 | 0 |

**Figure 1:**
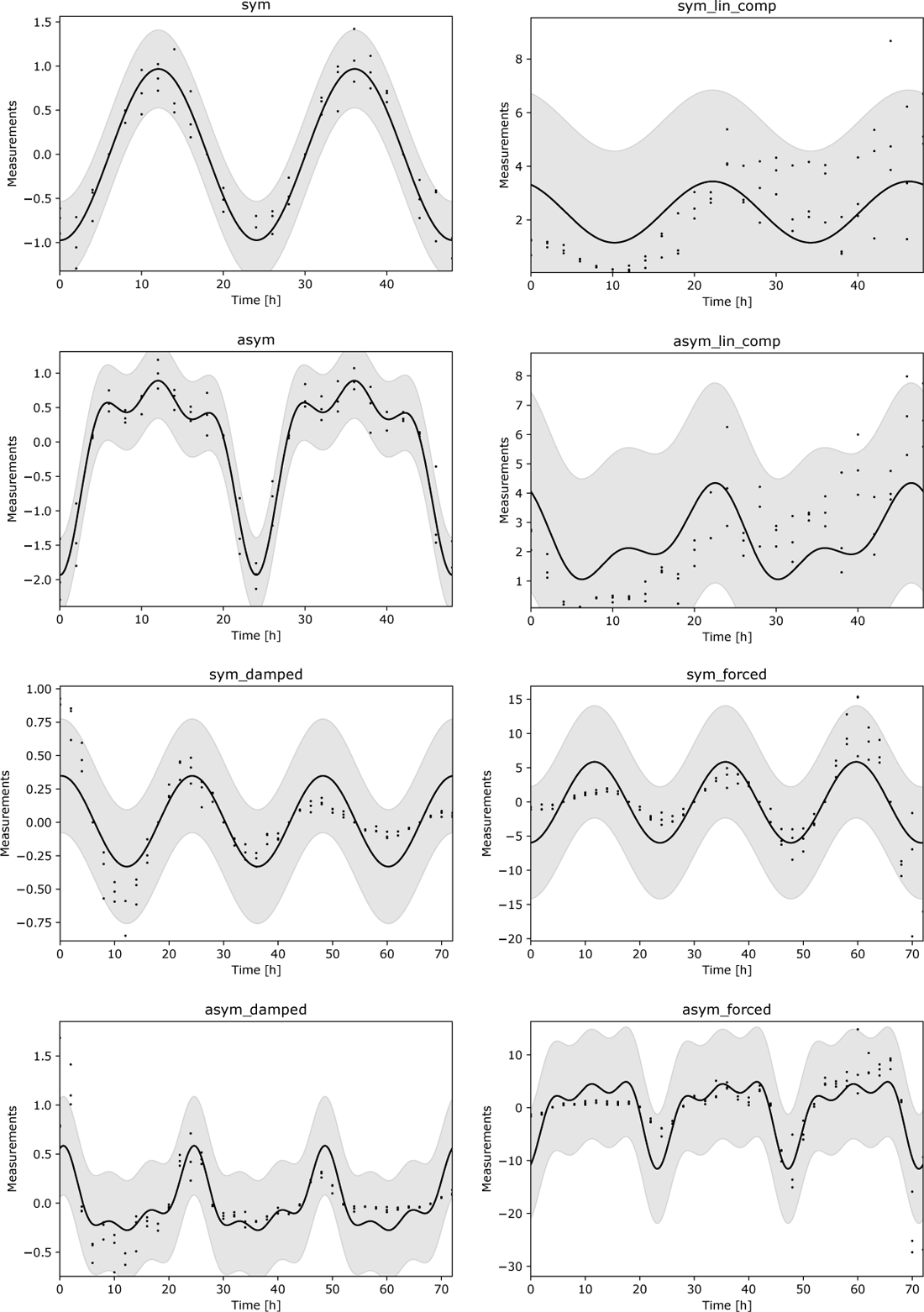
Results of the fitting process performed with a multi-component cosinor model. CosinorPy was used to identify the model that achieves the best fit for each scenario. The model is not able to account for forced/damped rhythms (`sym_damped`, `sym_forced`, `asym_damped`, `asym_forced`) and/or the accumulation of a linear component around which rhythmicity occurs (`sym_lin_comp`, `asym_lin_comp`)

We can generalise a single-component cosinor model to obtain a model in equation 2. We proceeded with our analysis using this model. Since it is based on non-linear regression, it is less robust than the basic cosinor model. However, the generalised model can account for additional dynamical aspects in the observed data, such as damped or forced rhythmicity and accumulation of the MESOR. The results of the fitting process using a single-component generalised cosinor model are visualised in Figure 3 and summarised in Table 2 (a complete statistic of the evaluated models is available as Supplementary Table 3).

**Table 2:** Results of the analysis of a single-component generalised cosinor model. Abbreviations and symbols: *A* – amplitude; *ϕ* – acrophase; amplif. – amplification; lin. – linear component; *q*(·) – false-discovery rate adjusted p-value

| test | $A$ | $q(A)$ | $\phi$ | $q(\phi)$ | amplif. | $q(\text{amplif.})$ | lin. | $q(\text{lin.})$ |
| --- | --- | --- | --- | --- | --- | --- | --- | --- |
| sym | 0.863 | **0.0** | 3.142 | **0.0** | 0.005 | 0.088 | 0.003 | 0.207 |
| sym_lin_comp | 1.071 | **0.004** | -0.174 | 0.483 | -0.001 | 0.910 | 0.093 | **0.0** |
| asym | 1.194 | **0.0** | -3.079 | **0.0** | -0.003 | 0.624 | 0.001 | 0.861 |
| asym_lin_comp | 1.306 | **0.001** | 0.016 | 0.932 | -0.012 | 0.433 | 0.103 | **0.0** |
| sym_damped | 0.927 | **0.0** | -0.017 | 0.599 | -0.037 | **0.0** | 0.0 | 0.675 |
| sym_forced | 0.897 | **0.0** | -3.118 | **0.0** | 0.041 | **0.0** | 0.009 | 0.675 |
| asym_damped | 1.127 | **0.0** | 0.148 | 0.044 | -0.050 | **0.0** | 0.0 | 0.675 |
| asym_forced | 0.282 | 0.062 | -3.142 | **0.0** | 0.061 | **0.0** | 0.015 | 0.675 |

This model was able to adequately describe damped and sustained rhythms as well as the accumulation of the MESOR. However, since it is based on a single-component cosinor, the model was unable to accurately describe the rhythmicity in the data reflecting asymmetric rhythmicity and/or multiple peaks per period (see Figure 2 and Figures `asym`, `asym_lin_comp`, `asym_damped`, `asym_forced`). Moreover, even though the data in the test `asym_forced` reflects oscillatory behaviour, the test of rhythmicity period being different from zero was not significant after the adjustment for multiple tests using the false discovery rate (*q* = 0.062, see Table 2).

**Figure 2:**
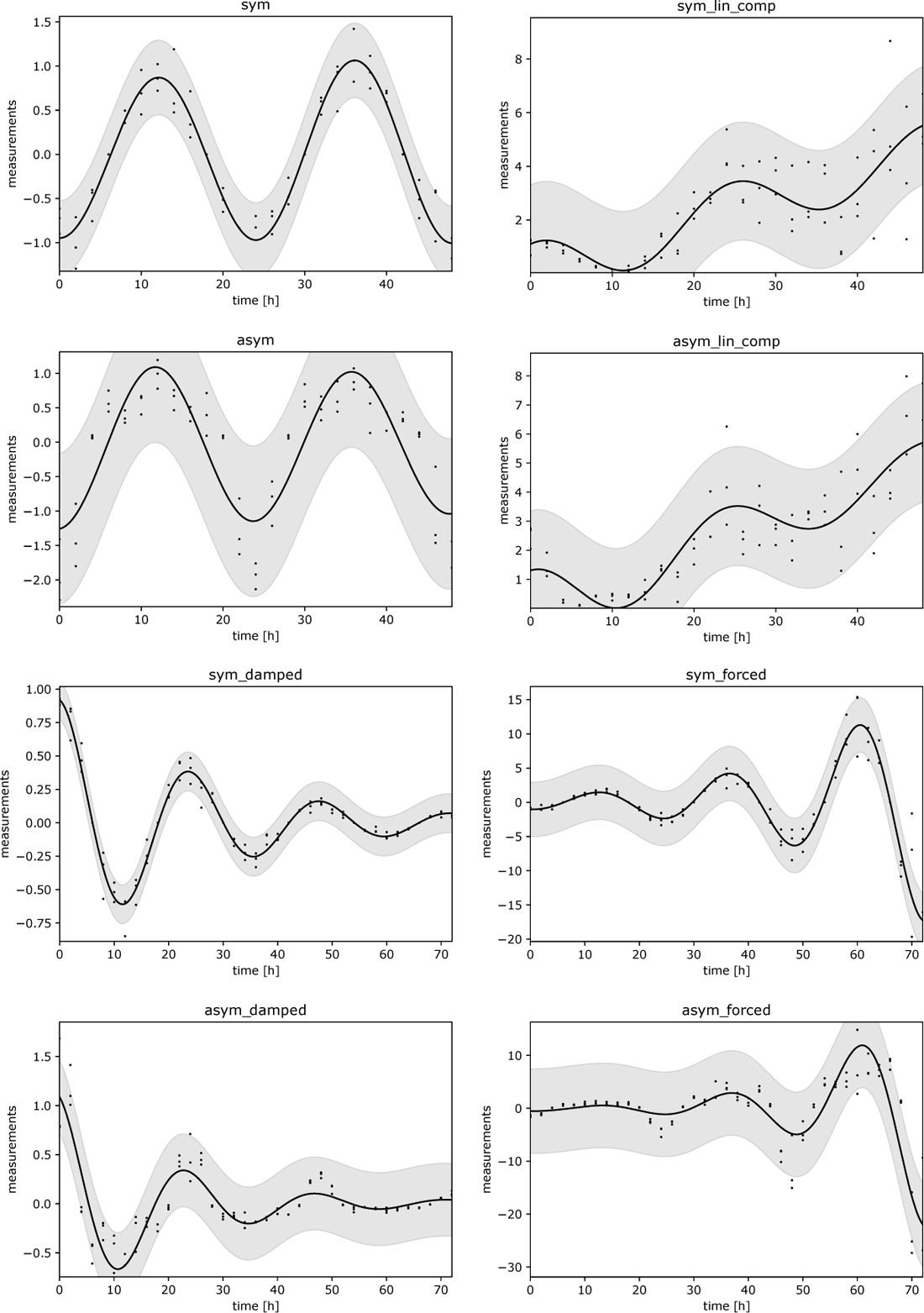
Results of the fitting process performed with a single-component generalised cosinor model. The model is not able to accurately describe the data reflecting asymmetric rhythmicity and/or multiple peaks per period (`asym`, `asym_lin_comp`, `asym_damped`, `asym_forced`). The model is able to account for forced/damped rhythms and/or the accumulation of a linear component around which rhythmicity occurs

**Figure 3:**
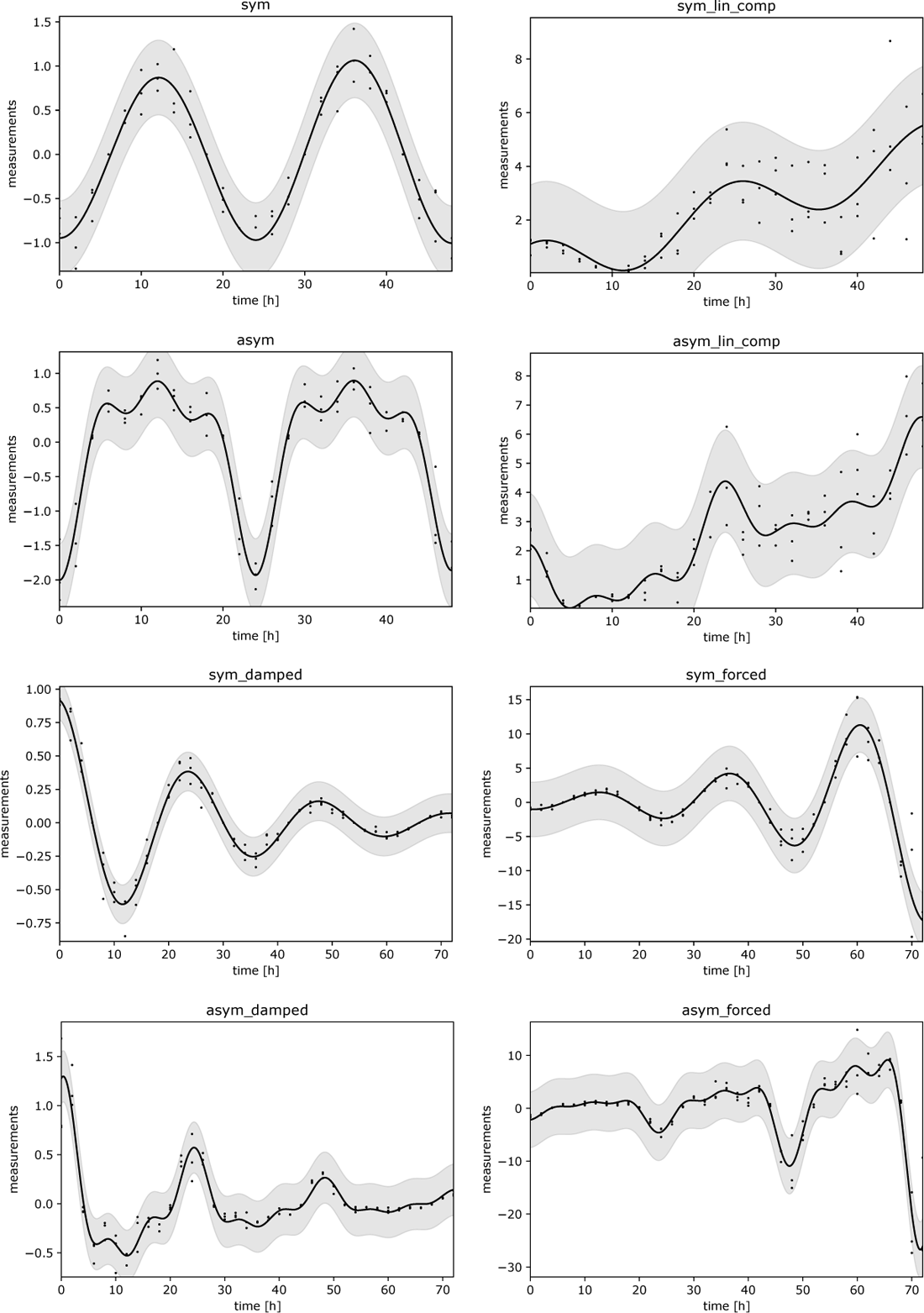
Results of the fitting process performed with a multi-component generalised cosinor model. CosinorPy was used to identify the model that achieves the best fit for each scenario. The model is able to describe the data reflecting asymmetric rhythmicity, multiple peaks per period, and is able to account for forced/damped rhythms and/or the accumulation of a linear component around which rhythmicity occurs

To address this problem, we applied a generalised multi-component cosinor model as described in Equation 4. CosinorPy was used to assess the optimal number of components in a cosinor model for each scenario. The optimal number of components equaled 1 for all symmetric cases and 3 for all asymmetric cases. The results of the fitting process are visualised in Figure 3, and the full statistics of the evaluated models are available as Supplementary Table 4. To identify the significance of the rhythmicity parameters, we ran the wrapper function that performs the bootstrap analysis of the obtained models. The results of this analysis are summarised in Table 3. As expected, the rhythmicity amplitudes were significant in all cases. Acrophase values were significant in all cases for which the acrophase values were set to *π*, namely `sym`, `asym_lin_comp`, `sym_forced`, and `asym_forced`. A complete statistics of the bootstrap analysis are available as Supplementary Table 5.

**Table 3:** Results of the bootstrap analysis of best-fitted generalised cosinor models. Abbreviations and symbols: *N* – number of components in a cosinor model, *A* – amplitude; *ϕ* – acrophase; amplif. – amplification; lin. – linear component; *q*(·) – false-discovery rate adjusted p-value

| test | $N$ | $A$ | $q(A)$ | $\phi$ | $q(\phi)$ | amplif. | $q(\text{amplif.})$ | lin. | $q(\text{lin.})$ |
| --- | --- | --- | --- | --- | --- | --- | --- | --- | --- |
| sym | 1 | 0.863 | **0.0** | 3.138 | **0.0** | 0.005 | 0.088 | 0.003 | 0.155 |
| sym_lin_comp | 1 | 1.071 | **0.0** | -0.176 | 0.415 | -0.001 | 0.910 | 0.093 | **0.0** |
| asym | 3 | 1.442 | **0.0** | 3.138 | **0.0** | -0.001 | 0.768 | 0.001 | 0.631 |
| asym_lin_comp | 3 | 1.588 | **0.0** | 0.094 | 0.357 | -0.005 | 0.609 | 0.102 | **0.0** |
| sym_damped | 1 | 0.927 | **0.0** | -0.013 | 0.487 | -0.037 | **0.0** | 0.000 | 0.631 |
| sym_forced | 1 | 0.897 | **0.0** | -3.120 | **0.0** | 0.041 | **0.0** | 0.009 | 0.473 |
| asym_damped | 3 | 1.054 | **0.0** | -0.138 | 0.128 | -0.034 | **0.0** | 0.001 | 0.155 |
| asym_forced | 3 | 1.408 | **0.0** | -2.981 | **0.0** | 0.037 | **0.0** | 0.017 | 0.318 |

### 3.2. Comparative analysis of rhythmic data

Using a basic or a generalised single- or multicomponent cosinor model, CosinorPy is able to evaluate the differential rhythmicity for an arbitrary pair of measurements. We applied a generalised cosinor model to analyse the differences between the pairs (`sym`, `sym_lin_comp`), (`asym`, `asym_lin_comp`), (`sym_damped`, `sym_forced`), and (`asym_damped`, `asym_forced`). Comparisons are visualised in Figure 4. Within each pair, differential rhythmicity was assessed using a wrapper function that performs a comparative bootstrap analysis. The results of this analysis are presented in Table 4 (complete statistics are available in Supplementary Table 6).

**Table 4:** Results of the comparative analysis. Abbreviations and symbols: *N*_1_, *N*_2_ – number of components in a cosinor model; *A* – amplitude; *ϕ* – acrophase; amplif. – amplification; lin. – linear component; *d*() – difference of parameter; *q*(·)–false-discovery rate adjusted p-value

| test | $N_1$ | $N_2$ | $d(A)$ | $q(d(A))$ | $d(\phi)$ | $q(d(\phi))$ | $d(\text{amplif.})$ | $q(d(\text{amplif.}))$ | $d(\text{lin.})$ | $q(d(\text{lin.}))$ |
| --- | --- | --- | --- | --- | --- | --- | --- | --- | --- | --- |
| sym vs. sym_lin_comp | 1.0 | 1.0 | 0.208 | 0.769 | 2.969 | **0.0** | -0.006 | 0.616 | 0.090 | **0.000** |
| asym vs. asym_lin_comp | 3.0 | 3.0 | 0.146 | 0.872 | -3.044 | **0.0** | -0.004 | 0.616 | 0.100 | **0.000** |
| sym_damped vs. sym_forced | 1.0 | 1.0 | -0.030 | 0.872 | -3.107 | **0.0** | 0.078 | **0.000** | 0.009 | 0.365 |
| asym_damped vs. asym_forced | 3.0 | 3.0 | 0.354 | 0.769 | -2.843 | **0.0** | 0.070 | **0.000** | 0.016 | 0.309 |

**Figure 4:**
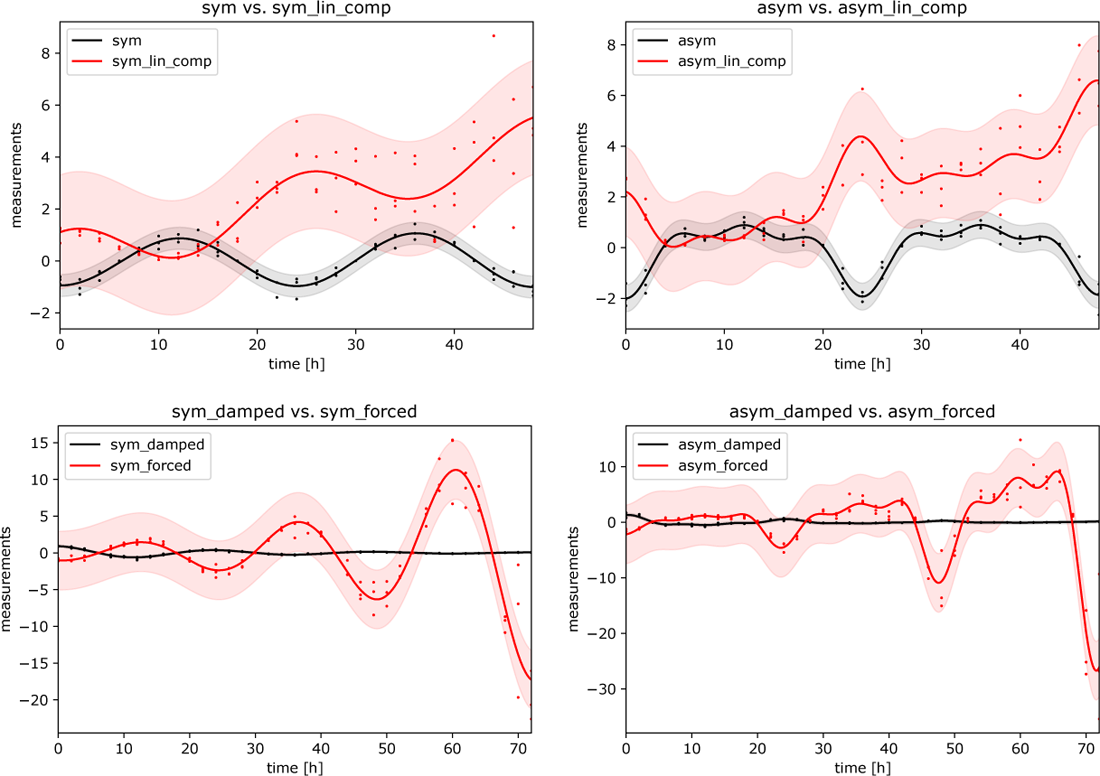
Results of the comparison analysis using a multi-component generalised cosinor model. The models that achieve the best fit for each scenario and identified in the previous steps were used in the analysis

### 3.3. Evaluating errors of different cosinor models

To rigorously assess the errors produced by different cosinor models, we generated a set of rhythmic data reflecting different types of oscillatory behaviour. That is, we generated samples reflecting (1) symmetric and asymmetric rhythmicity with single or multiple peaks per period, and (2) samples reflecting damped, sustained, and forced rhythmicity. For each of the combinations of (1) and (2), 100 samples were generated. Moreover, each of these samples was combined with different levels of noise, which were set to 10%, 20%, 50%, and 100% of the generated data. We used the generated samples in a combination with different cosinor models, namely a single-component cosinor model (C1), a single-component generalised cosinor model (C1G), a 3-component cosinor model (C3), and a 3-component generalised cosinor model (C3G). We evaluated the mean squared error (MSE) of evaluated rhythmicity amplitudes and acrophases for each of these models. MSE of evaluated rhythmicity amplitudes are presented in Table 5, and MSE of the evaluated rhythmicity acrophases are presented in Table 6. The distribution of errors for noise levels of 10%, 20% and 50% are visualised in Figures 5 and 6. The entire analysis is available as an interactive Python Jupyter notebook at https://github.com/mmoskon/CosinorPy/blob/master/analyse_errors.ipynb.

**Table 5:**
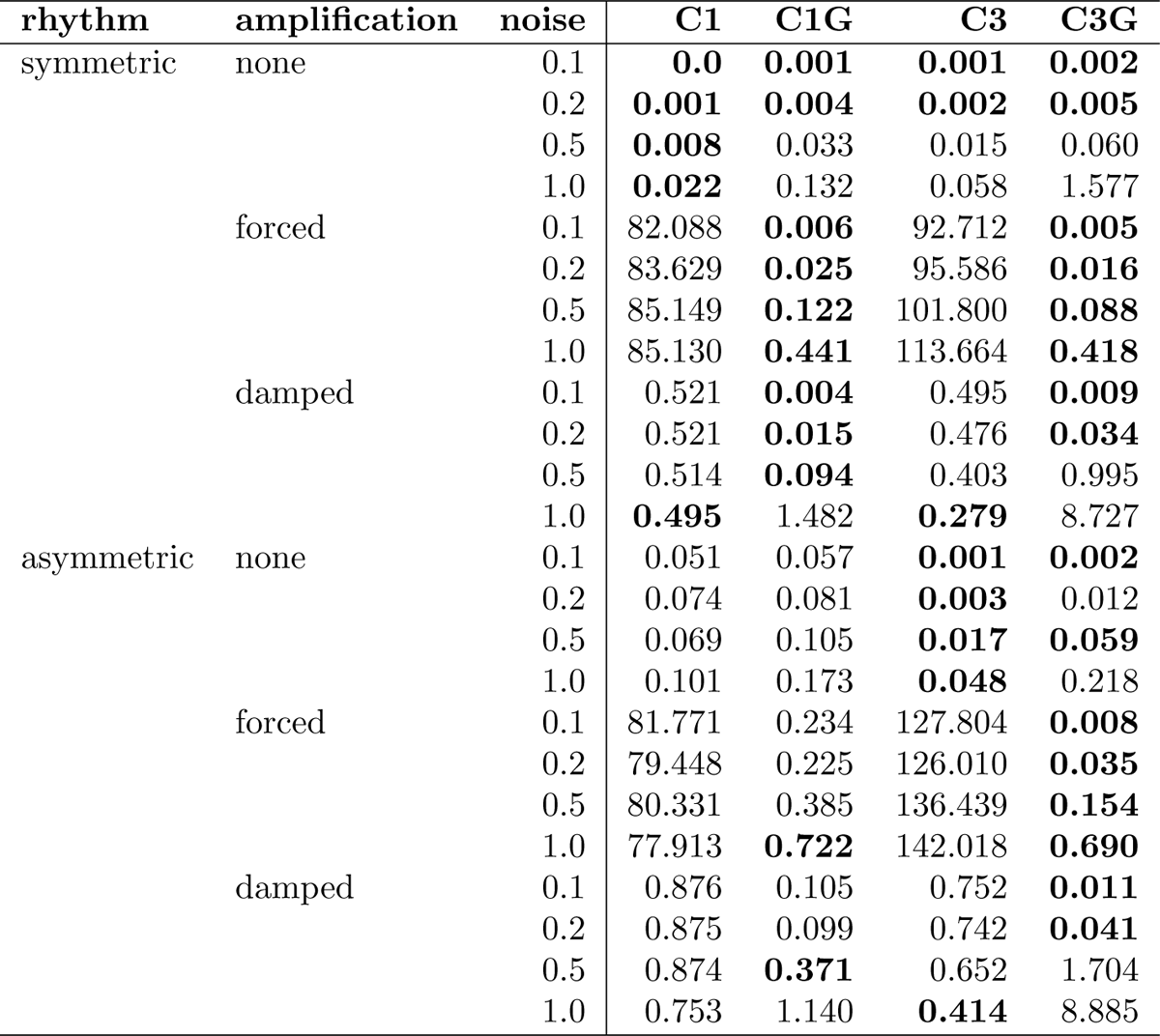
Mean squared errors of the evaluated rhythmicity amplitudes. Abbreviations: C1 – single-component cosinor model; C3 – 3-component cosinor model; C1G – generalised single-component cosinor model; C3G – generalised 3-component cosinor model

**Table 6:** Mean squared errors of evaluated rhythmicity acrophases. Abbreviations: C1 – single-component cosinor model; C3 – 3-component cosinor model; C1G – generalised single-component cosinor model; C3G – generalised 3-component cosinor model

| rhythm | amplification | noise | C1 | C1G | C3 | C3G |
| --- | --- | --- | --- | --- | --- | --- |
| symmetric | none | 0.1 | **0.0** | **0.0** | 0.006 | 0.006 |
|  |  | 0.2 | **0.001** | **0.001** | 0.027 | 0.028 |
|  |  | 0.5 | **0.006** | **0.006** | 0.132 | 0.126 |
|  |  | 1.0 | **0.031** | **0.030** | 0.248 | 0.228 |
|  | forced | 0.1 | **0.004** | **0.0** | 0.120 | **0.004** |
|  |  | 0.2 | **0.004** | **0.001** | 0.120 | 0.017 |
|  |  | 0.5 | **0.007** | **0.004** | 0.147 | 0.111 |
|  |  | 1.0 | **0.013** | **0.015** | 0.218 | 0.271 |
|  | damped | 0.1 | **0.005** | **0.001** | 0.122 | 0.031 |
|  |  | 0.2 | 0.017 | **0.007** | 0.187 | 0.121 |
|  |  | 0.5 | 0.079 | **0.046** | 0.457 | 0.384 |
|  |  | 1.0 | **0.354** | **0.361** | 0.798 | 0.863 |
| asymmetric | none | 0.1 | 0.227 | 0.228 | **0.043** | **0.042** |
|  |  | 0.2 | 0.274 | 0.275 | **0.122** | **0.154** |
|  |  | 0.5 | 0.226 | 0.225 | **0.182** | **0.181** |
|  |  | 1.0 | **0.287** | **0.287** | 0.508 | 0.522 |
|  | forced | 0.1 | 0.214 | 0.209 | 0.320 | **0.043** |
|  |  | 0.2 | 0.200 | 0.211 | 0.253 | **0.032** |
|  |  | 0.5 | **0.223** | 0.241 | 0.283 | **0.208** |
|  |  | 1.0 | **0.241** | **0.265** | 0.324 | 0.541 |
|  | damped | 0.1 | 0.209 | 0.215 | 0.277 | **0.088** |
|  |  | 0.2 | 0.252 | 0.257 | **0.204** | **0.220** |
|  |  | 0.5 | **0.240** | **0.258** | 0.386 | 0.579 |
|  |  | 1.0 | **0.582** | **0.549** | 1.055 | 1.071 |

**Figure 5:**
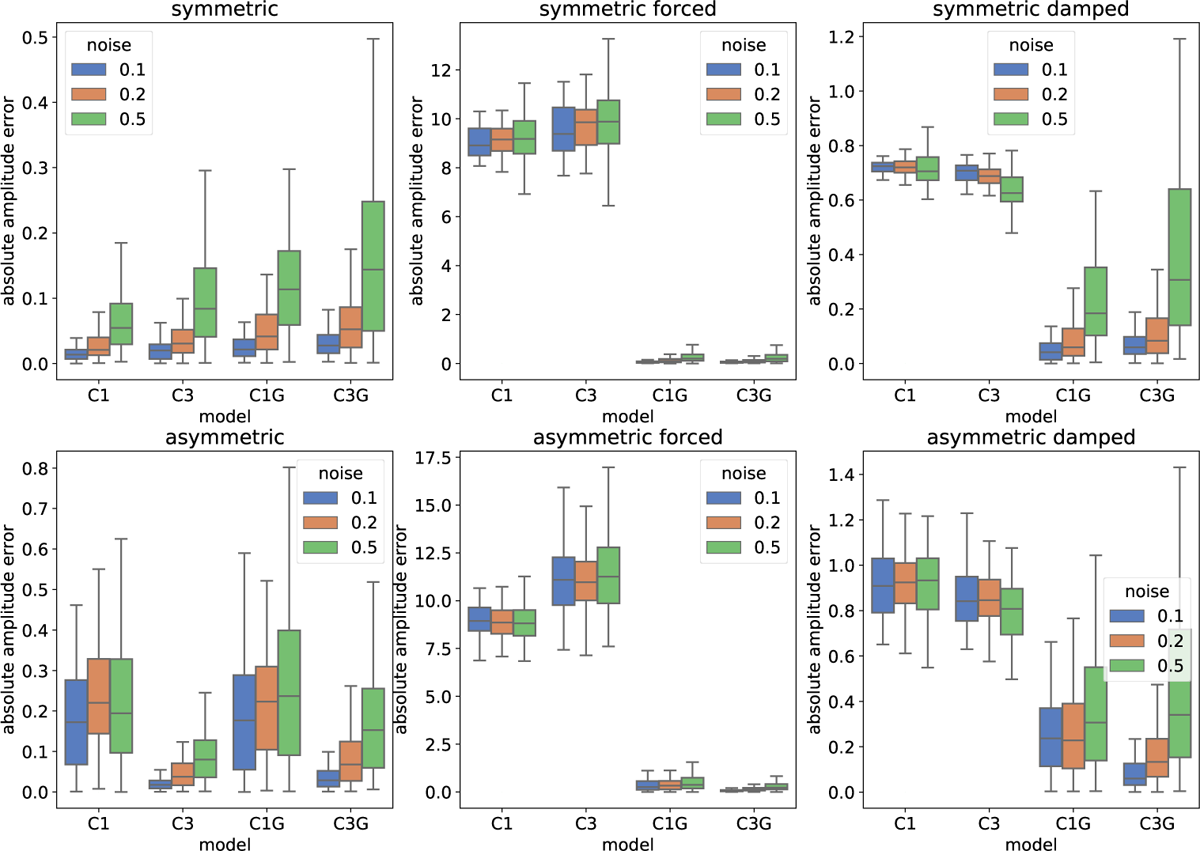
Distributions of amplitude errors for different cosinor models using different types of rhythms and different levels of noise. Abbreviations: C1 – single-component cosinor model; C3 – 3-component cosinor model; C1G – generalised single-component cosinor model; C3G – generalised 3-component cosinor model

**Figure 6:**
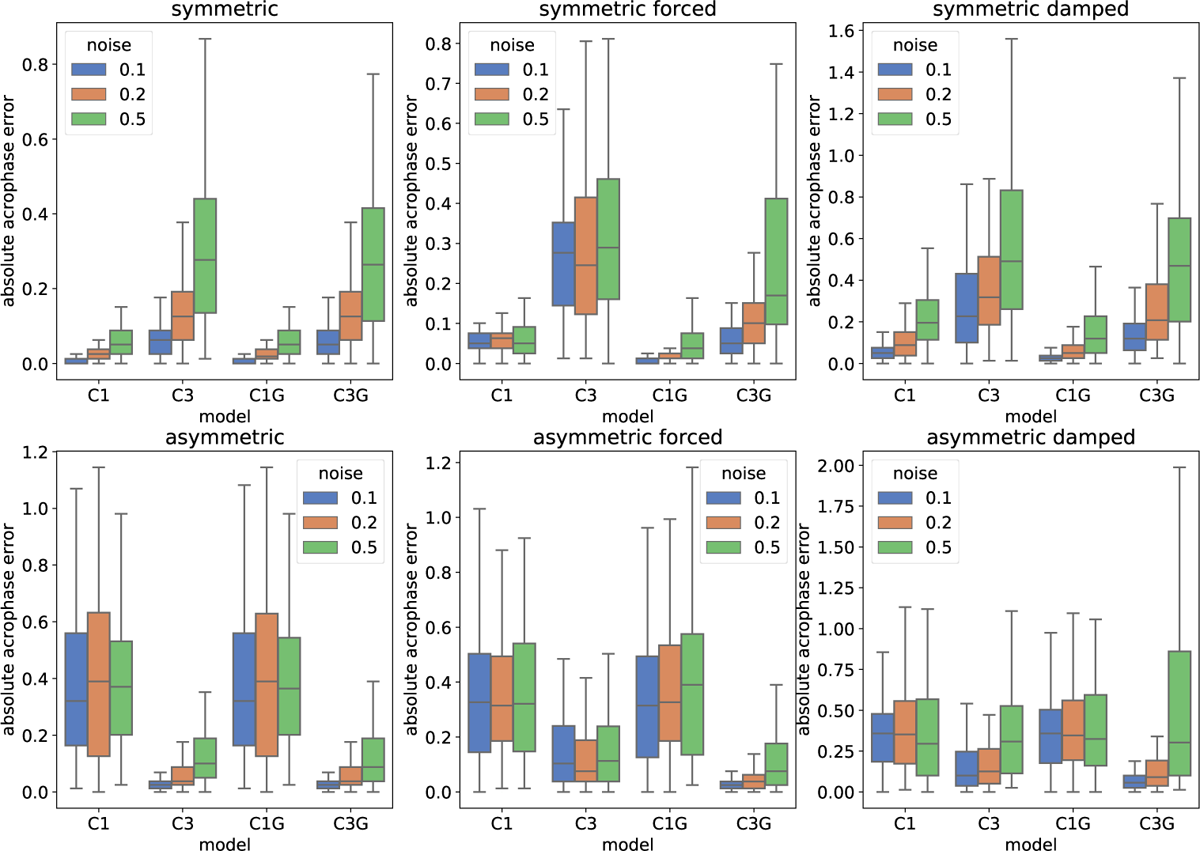
Distributions of acrophase errors for different cosinor models using different types of rhythms and different levels of noise. Abbreviations: C1 – single-component cosinor model; C3 – 3-component cosinor model; C1G – generalised single-component cosinor model; C3G – generalised 3-component cosinor model

The results indicate that the proposed generalised cosinor models can accurately reproduce the oscillatory signal even when rhythms are damped or forced. As already shown in our previous analyses, asymmetric rhythms are better reproduced with a multi-component cosinor model. However, when the noise is increased above 50% of the signal, it is better to use a model with a smaller number of components to reduce the error, especially if rhythms are damped. This derives from the fact that a model with a larger number of components can easily be over-fitted to the noisy data. When noise becomes prevalent, simpler models tend to perform better and are more robust. This problem can be controlled by selecting the most suitable model for each dataset separately. That is, we should opt to select the model that is capable of describing the dataset with the smallest number of components. This can be performed automatically using the functionalities provided by CosinorPy [9].

## 4. Conclusions

The extended version of the CosinorPy package provides a suite of methods that can be used in the analysis of different types of rhythmic data. Most importantly, it can be used to assess the statistics of the rhythmicity parameters even when the observed data reflect multiple asymmetric peaks per one period of a rhythm. Moreover, the extended package provides an implementation of generalised single- and multicomponent cosinor models, which can describe damped or forced rhythms as well as the accumulation of the MESOR. The statistics of assessed fits can be evaluated even with these relatively complex models using bootstrapping of confidence intervals and p-values. We believe that the proposed extensions of CosinorPy will find even wider applicability than its original implementation.

## Acknowledgements

I would like to thank all the past, current and future users of CosinorPy for using the package and for providing the critical feedback, which allows the package to progress further.

